# ROCK inhibitor Ripasudil reduces Alzheimer’s pathological proteins and enhances cellular bioenergetics

**DOI:** 10.64898/2026.11.02.761117

**Authors:** Yuehan Li, Zhiqing Liu, Xavier A. Cambi, Jane Paterson, Gloria A. Benavides, Xiaosen Ouyang, Alec Candib, Joseph Zaia, Victor Darley-Usmar, Chao Zheng, Jianhua Zhang, Weiming Xia

**Affiliations:** Department of Pharmacology, Physiology & Biophysics, Boston University Chobanian and Avedisian School of Medicine, Boston, MA 02118, USA; Geriatric Research Education and Clinical Center, Bedford VA Healthcare System, Bedford, MA 01730, USA; Azrieli Centre for Neuro-Radiochemistry, Brain Health Imaging Centre, CAMH, Departments of Psychiatry, Chemistry, Pharmacology and Toxicology, University of Toronto, Toronto, Ontario, M5T 1R8 Canada; Department of Pathology, University of Alabama at Birmingham, Birmingham, AL 35294, USA; Department of Biochemistry, Boston University Chobanian and Avedisian School of Medicine, Boston, MA 02118, USA; Department of Biological Science, Kennedy College of Sciences, University of Massachusetts Lowell, Lowell, MA 01854, USA

**Author notes:** CORRESPONDING AUTHOR: Weiming Xia, PhD, Geriatric Research Education and Clinical Center, Bedford VA Healthcare System, 200 Springs Road, Bedford, MA, USA.

**Keywords:** Alzheimer’s disease, ripasudil, ROCK, therapeutics, iPSC, proteomics, mitochondria

## Abstract

Rho-associated kinase (ROCK) inhibitors have been explored as an Alzheimer’s Disease (AD) therapeutic agent. In this study, we aim to determine the efficacy of a potent ROCK inhibitor, ripasudil, in impacting AD pathological protein tau in mice. The time-course study showed that ripasudil rapidly penetrated mouse brains, reaching maximal concentration half an hour post-injection. To determine the efficacy of ripasudil, we injected *App^NL-G-F^* knock-in mice with ripasudil at 30 mg/kg and sacrificed them after three hours. We used Tandem Mass Tag labels and compared the proteomic profiles of mouse brains exposed to ripasudil to those exposed to saline. We also treated AD-patient-derived Neural Stem Cells (NSCs) with ripasudil and performed a Seahorse Mito Stress Test to evaluate their impact on mitochondrial activity. Bioinformatics analysis of the proteomic profiles revealed differentially enriched components of chromosome. The Mito Stress Test of NSCs revealed significant increases in basal oxygen consumption, ATP-linked respiration, maximal respiration, and reserve capacity oxygen consumption compared to untreated NSCs. Importantly, we found a significant reduction of Tau and pTau 396 in brains of mice treated with ripasudil. Overall, our findings highlight the feasibility and promise of ROCK inhibitor ripasudil as an AD therapeutic agent.

## INTRODUCTION

Alzheimer’s Disease (AD) is a progressive and irreversible neurodegenerative disease. AD is characterized by extracellular β-amyloid (Aβ) containing neuritic plaques and intracellular phosphorylated tau (pTau) containing neurofibrillary tangles (NFTs) ^1^. AD is the leading cause of dementia, causing progressive loss of neurons and synapses, especially in the cerebral cortex and hippocampus. Neuroinflammation, astrocytosis, and the activation of microglia are also observed in brains of AD patients. Functional deterioration in AD is accompanied by progressive neuronal loss. There are various mechanisms that can impair neuronal function, one of which is mitochondrial dysfunction ^2^. Mitochondria are involved in processes such as cell death, apoptosis, and reactive oxygen species ^3^ production that are all dysregulated in neurodegenerative diseases such as AD ^4^. AD causes severe cognitive impairment and decline in executive functions ^5^, and more than six million Americans are living with AD, and by 2060, this number is projected to rise to nearly 14 million ^6^.

Until five years ago, the U.S. Food and Drug Administration (FDA) approved only symptomatic treatments for AD, rather than disease-modifying ones ^7^. This is the case despite the fact that research has identified familial mutations in several genes that directly cause AD, such as the genes that encode the amyloid precursor protein (APP) and the presenilin 1 and 2. Therapeutic development in the past three decades has been mainly focused on Aβ, which is derived from β- and γ-secretase cleavages of amyloid precursor protein (APP). Recently, FDA approved disease-modifying therapeutics that directly target Aβ, and these immunotherapies using humanized monoclonal antibodies have shown clinical efficacy ^8, 9^. The antibodies Lecanemab and Donanemab have been prescribed for patients with mild cognitive impairment (MCI) and mild AD ^10, 11^. However, these drugs come with amyloid-related imaging abnormalities (ARIA), which can lead to edemas and brain hemorrhages ^3^. Therefore, there is a huge unmet need for safe, efficacious, disease-modifying therapeutics for AD.

One direction is to explore Rho-associated kinase (ROCK) inhibitors. ROCKs are serine/threonine kinases that are the downstream targets of the small GTPase Rho. Rho proteins can be activated by guanine nucleotide exchange factors (GEFs), which are themselves activated by receptor tyrosine kinases (RTKs), G protein-coupled receptors (GPCRs), cytokines, and integrins. Rho-GTP then activates ROCK1 and ROCK2, which have a broad range of substrates and are responsible for diverse cellular responses, including actin cytoskeleton organization, cell adhesion and motility, proliferation and apoptosis, remodeling of the extracellular matrix, and smooth muscle cell contraction ^12^. Two homologs ROCK1 and ROCK2 share 65% of their overall amino acid sequences, and 92% of their kinase domains are identical. While both isoforms are ubiquitously expressed, ROCK2 has an elevated expression in the brain ^13^. In the brain, ROCKs are involved in elongating neurites, facilitating neuronal architecture, regulating blood-brain barrier (BBB) integrity, and maintaining neuronal processes that underlie synaptic transmission and cognitive functions ^13, 14^.

Overactivation of ROCKs leads to inflammation, oxidative stress, neuronal loss, and impaired synaptic transmission ^15^. Therefore, pharmacological inhibition of ROCKs has shown potential of neuroprotective effects in AD ^16^ and other neurological diseases, such as Parkinson’s disease (PD) ^17^, multiple sclerosis (MS) ^18^, and amyotrophic lateral sclerosis (ALS) ^19^. The only ROCK inhibitor in clinical trial for AD is fasudil, which is clinically approved and used for the treatment of cerebral vasospasm in Japan and China ^20^. Fasudil has been shown to decrease neuronal loss and increase neurological function in animal models of ischemic stroke ^21^, both improving learning and memory in rats ^21^ and improving memory and mental function in patients with cerebrovascular dementia ^22^. Furthermore, fasudil has been shown to promote lengthening of dendritic arbors in APP/PS1 transgenic mice and protect against Aβ-induced hippocampal neurodegeneration in rats by suppressing inflammatory responses ^23, 24^.

Pharmacological inhibition of ROCKs decreases tau phosphorylation in cellular and *in vivo* models of tauopathy ^14, 25–28^. The dysregulation of microtubule-associated protein tau phosphorylation is a pathological hallmark of AD and a precursor of NFTs ^29^. Tau is phosphorylated by ROCKs at several sites, and *in vitro* studies using the neuroblastoma cell line N2a showed that ROCK inhibition reduces aggregated tau levels by activating cellular protein degradation machinery ^30^. Fasudil can decrease tau phosphorylation in primary cultures of mouse hippocampal neurons ^31^. *In vivo* studies with APP/PS1 transgenic mice have shown that pharmacological inhibition of ROCK can inhibit the formation of pTau, a modified form of the tau protein that detaches from neuronal microtubules and leads to instability, aggregation, and the formation of NFTs ^32^. Furthermore, in an *in vivo* study with rTG4510 tau transgenic mice, ROCK inhibitors reduced pTau and oligomeric tau levels, promoted tau degradation by inhibiting the activity of the tau kinases GSK3β, Cdk5, and caspase, and activated tau phosphatase PP2A and proteasome pathways ^33^. Pharmacological inhibition of ROCKs induced autophagy in pathways targeting both soluble and aggregated forms of tau, the same mechanism as that degrading Aβ ^25^. Recent studies have also reported that ROCK inhibition in neurons reduces tau mRNA ^16^. In Tau transgenic PS19 mouse model exposed to fasudil, a decrease in mitochondrial complex IV subunit I was found in the striatum and thalamic regions; chronic dosing of fasudil for 2 weeks resulted in significantly decreased mitochondrial outer membrane and electron transport chain (ETC) protein abundance, as well as ETC activities ^34^.

Ripasudil is a ROCK inhibitor with a higher potency for ROCK2 than ROCK1, compared to fasudil ^35^. Fasudil inhibits ROCK1/2 with a poor IC_50_ (740 nM), which is difficult to achieve *in vivo* particularly in the brain^36^. In contrast, ripasudil is a much more potent ROCK1/2 inhibitor, with an IC_50_ of 51 and 19 nM, respectively^36, 37^. Fasudil is an isoquinoline derivative, and ripasudil is a chemically-modified ROCK inhibitor with a fluorine atom at the C4 position of the isoquinoline ring ^38^. Ripasudil has been approved in Japan for treating glaucoma and ocular hypertension since 2014 ^39^. Given the neuroprotective effects of fasudil against tau phosphorylation, we aimed to evaluate in vivo output to measure the efficacy of the potent ROCK inhibitor ripasudil.

## METHODS

### Drug treatment

Wild-type mice (C57BL/6J, Jackson Lab, Bar Harbor, ME) were used to determine the time course of ripasudil (Selleck Chemicals LLC, Houston, TX). Saline or drug were administered by intraperitoneal (i.p.) injection at 30 mg/kg, and brains and blood were harvested at 0.5, 1, 2, 4, 8 and 24 hours to determine the drug exposure. The brains were collected and stored at -80°C until further processing. All studies were approved by the institutional IACUC at the Bedford VA Healthcare System.

### *App^NL-G-F^* knock-in mice

*App^NL-G-F^* knock-in mice do not overexpress APP but carry 3 AD-linked familial mutations: APP KM670/671NL (Swedish), APP I716F (Iberian), and APP E693G (Arctic). It has been shown that the Swedish mutation increases total Aβ production, the Iberian mutation increases the Aβ42/Aβ40 ratio, and the Arctic mutation increases oligomerization while decreasing proteolytic degradation of Aβ^40, 41^. Five-month-old mice were used for efficacy studies by i.p. dosing of ripasudil. To determine the efficacy of ripasudil, animals were dosed at 30 mg/kg and euthanized by CO_2_ inhalation after three hours. The brains were collected and stored at -80°C until further processing.

### Quantification of brain Tau and pTau 396 levels

ELISA was performed to quantify brain levels of Tau and pTau 396. A lysis buffer (2% SDS, 0.5M TEAB, protease and phosphatase inhibitor cocktail) was added to brain tissue at a 5:1 ratio (0.2 g/mL). Tissue was homogenized by TissueLyser LT (Qiagen, Valencia, CA). Tissue homogenates were centrifuged at 16,000 x g for 20 minutes at 4°C and loaded onto ELISA plates coated with the corresponding primary antibodies. Plates were read using the MSD Sector Imager 2400 (Rockville, MD). The same lysates were used for proteomic profiling ^42^.

### Determination of brain levels of Ripasudil

The established LC-MS/MS method was adapted to determine the levels of ripasudil in the brains of mice given a single dose of ripasudil by i.p. injection ^28^. An UltiMate 3000 UHPLC automated system coupled with a TSQ Quantiva triple quadrupole Mass Spectrometer (Thermo Fisher, Waltham, MA) was used for these experiments. A pre-calculated volume (based on 0.2 g/mL, ratio 1:5, w/v) of ice-cold water-methanol (30:70, v/v) was added to each brain sample, followed by homogenization using a TissueLyser LT (Qiagen, Valencia, CA). The brain homogenates were centrifuged at 17,000 × g for 20 minutes at 4°C, and 50 μL of the supernatant was mixed with 130 μL of ice-cold acetonitrile and 20 μL of an internal standard solution. Samples were vortexed vigorously in ice between the steps throughout the process. Following mixing, the samples were centrifuged at 20,000 × g for 5 minutes at 4°C. The final supernatants were transferred to HPLC vials for LC-MS/MS analysis.

An ACQUITY UPLC BEH C18 column (75 × 2.1 mm, 1.7 μm particle size; Waters, Milford, MA) was used to achieve chromatographic separation, with the mobile phases consisted of water with 0.1% formic acid (mobile phase A) and acetonitrile with 0.1% formic acid (mobile phase B). A linear gradient elution, running from 3% to 95% B over nine minutes, was employed, maintaining at 95% B for ten minutes, returning to 3% B in one minute, and equilibrating for five minutes at 3% B, at the flow rate of 0.10 mL/min.

Mass spectrometric detection was performed using a heated electrospray ionization (H-ESI) source operating in positive ion mode. Data was acquired in selected reaction monitoring (SRM) mode, with the ion spray voltage set to 3.7 kV, and the ion transfer tube temperature maintained at 325°C. The monitored mass-to-charge (m/z) transitions were 324 → 146 for ripasudil and 315 → 176 for internal standard, with a second transition for each analyte used for confirmation.

### Preparation and differential labelling of proteomics samples

The established proteomics method was adapted to label the samples ^43^. Brain tissue was homogenized by vortexing in a sample lysis buffer with 2% SDS, 0.5M tetraethylammonium bicarbonate (TEAB), and phosphatase/protease inhibitor cocktails. Homogenates were centrifuged at 17,000 x g for 20 minutes at 4⁰C, and the supernatants were labeled with Tandem Mass Tag (TMT) 10plex Isobaric Labeling Reagents (ThermoFisher). In brief, 100 µg of protein lysates were reduced with Tris(2-carboxyethyl)phosphine (TCEP) at 37⁰C for one hour, followed by alkylation with iodoacetamide in the dark for 0.5 hour. Four parts methanol was added to each sample, followed by vortexing and the addition of one part chloroform, and finally additional vortexing was performed for methanol-chloroform precipitation. To achieve phase separation, three parts of water were added, and the samples were vigorously vortexed and underwent centrifugation at 14,000 x g for four minutes at room temperature. The organic phase containing protein precipitates at the interface, and the samples were washed twice with four parts methanol. The air-dried precipitates were then re-suspended in 100 µl of 50 mM TEAB and digested with trypsin overnight at 37⁰C. For the TMT labeling, the trypsin-digested peptides were incubated with the TMT 10plex reagent (40µl in anhydrous acetonitrile) for one hour at room temperature. The reaction was then quenched by adding 8 µl of 5% hydroxylamine solution. The combined TMT-labeled peptides were dried under a SpeedVac, reconstituted in a diluted trifluoroacetic acid (TFA) solution, and cleaned using an Oasis HLB 96-well µElution plate (Waters).

### LC-MS/MS analysis

The established proteomics method was adapted for LC-MS/MS analysis ^43^. In brief, LC-MS/MS analysis was conducted using a Q Exactive Orbitrap Mass Spectrometer (Thermo Fisher Scientific) coupled with a Dionex Ultimate 3000 RSLCnano system equipped with an EASY-Spray ion source. The TMT-labeled peptides were separated on a C18 reverse-phase nano column (Acclaim PepMap, 75 µm x 500 mm, Thermo Fisher) with a linear gradient of 2–35% acetonitrile in 0.1% formic acid (FA), at a constant flow rate of 300 nL/min over 3 hours. The Q Exactive was operated in a positive-ion mode, with the ESI spray voltage set at 1.6 kV. A full scan MS spectrum (300–1800 m/z) was acquired at a mass resolution of 70,000, with an automatic gain control (AGC) target of 3×10^6^. The 12 most intense peptide ions from each scan were selected for higher-energy collision-induced dissociation (HCD)-MS/MS analysis (normalized collision energy 32) in the Orbitrap at a mass resolution of 35,000 and an AGC value of 1×10^5^. The maximum fill times were 100 milliseconds for full scans and 140 milliseconds for HCD-MS/MS scans. The ions with unassigned charge states, singly charged species, and those with more than seven charges were excluded. Dynamic exclusion was set to 20 seconds, with a relative mass window of 10 ppm. Data was acquired using the Thermo Xcalibur 3.0.63.

### Data processing and protein identification/quantification

The established proteomics method was adapted for data processing and protein identification ^43^. In brief, raw data was processed using a Proteome Discoverer (Version 2.1, Thermo Fisher Scientific) and searched against the mouse Universal Protein Resource sequence database using the following parameters: carbamidomethylation of cysteine as a fixed modification; trypsin digestion allowing up to two missed cleavages; oxidation of methionine, and TMT 10plex labeling at the N-terminus and lysine as variable modifications; MS tolerance set to 10 ppm; MS/MS tolerance set to 0.02 Da; and a false discovery rate (FDR) of less than 0.01 at both peptide and protein levels. A minimum peptide length of five amino acids was required. Protein grouping was enabled, such that if one protein sequence was equal to or entirely contained within the set of peptides of another protein, these proteins were grouped together.

### Bioinformatic analysis

Quantified proteins were analyzed in RStudio (Posit Software, PBS, Boston, MA). Protein abundances were log-transformed before differential expression analysis with limma-trend ^44^. Ripasudil-treated males were compared to saline-treated males, and ripasudil-treated females were compared to saline-treated females. Pathway enrichment analysis was performed with fgsea ^45^, with proteins ranked by t value and pathways sourced from MSigDB Gene Ontology gene sets ^46–48^. Statistical significance was defined as FDR<0.05 after BH correction ^49^. Visualizations were created with ggplot2 ^50^. Network graphs were created with igraph ^51^ and visualized with ggraph ^52^.

### Cell culture

Induced pluripotent stem-cell-derived Neural Stem Cells (NSCs) were used for cell culture. Those cells were well characterized and extensively utilized in our previous studies ^53–55^. We tested ripasudil in NSCs and established it as a drug screening platform with potential for scalability for high-throughput studies. Tissue culture flasks were first coated with Poly-L-Ornithine (PLO) at 15 µg/mL. After incubation at room temperature for two hours or at 2–8°C overnight, PLO was aspirated, and the flask was then coated with Laminin at 10 µg/mL. The same incubation procedure was followed, and NSCs were seeded accordingly. NPCs were grown in the STEMdiff Neural Progenitor medium. The night before the mitochondrial function experiment, NPCs were seeded in an XF Pro PDL cell culture microplate. The cultures were kept in a humid incubator (5% CO_2_, 37 ℃).

### Determination of mitochondrial function

To measure mitochondrial function in NSCs, the Seahorse XF Pro Analyzer (XF Pro) was used. The XF Pro measures and reports the oxygen consumption rate (OCR) of live cells in a 96-well format. The optimal seeding density of the cells was established. In all experiments, a seeding density of 40,000 cells per well was used. Cells were plated the day before the assay. After one hour of stabilization, cells were treated with either fresh media or ripasudil at 10 µM overnight. On the day of assay, cells were incubated in an XF Assay Medium (Seahorse XF DMEM, 10mM glucose, 1mM pyruvate, 2mM glutamine) with and without ripasudil at 10 µM for 1 hr. Then mitochondrial function was continuously measured for two hours, at baseline and after the sequential injection of oligomycin, carbonyl cyanide 4-(trifluoromethoxy)phenylhydrazone (FCCP), and antimycin A/Rotenone. The concentrations used were 1.5 µM, 2 µM, and 0.5 µM, respectively. All compounds were prepared fresh with an XF Assay Medium on the morning of the experiment. Ten wells of control and six wells of ripasudil-treated wells were used for the assay. Analysis was performed on Wave Pro software (Agilent, 10.2.0).

### [^18^F] BCPP-EF mitochondrial complex 1 function (MC-1) positron emission tomography (PET)

[^18^F]BCPP-EF was synthesized with high radiochemical purity of 99.83% and a molar activity (*A*_m_) of 4.38 Ci/µmol. Dynamic PET imaging was performed following bolus injection of [^18^F]BCPP-EF, with image acquisition conducted over 90 min after radiotracer administration. Six-month-old female wild-type C57BL/6 mice received either saline or ripasudil at 30 mg/kg by intraperitoneal (*i.p*.) injection 2 hours prior to PET imaging. PET image processing and quantification were performed according to previously published protocols ^56^. Briefly, dynamic PET data were used to generate whole-brain standardized uptake value time–activity curves to assess tracer uptake kinetics. Cerebellum-normalized standardized uptake value ratio (SUVR_CB_) images were subsequently generated, using the cerebellum as a pseudo-reference region to normalize brain uptake. These analyses were used to assess [^18^F]BCPP-EF uptake as an *in vivo* PET measure related to mitochondrial complex 1 availability.

### Statistical analysis

Data analysis and graph processing was performed using Prism 10 (GraphPad Software LLC, San Diego, CA). Comparisons between two groups were performed with unpaired t-tests. Two-way ANOVA was applied to identify significant variance in App^NL-G-F^ knock-in mice. Following significant effects, Tukey’s post-hoc tests were performed. The significance level (α) was set at 0.05.

## RESULTS

### Pharmacokinetics of ripasudil in mouse brains

To establish the pharmacokinetic profile of ripasudil, we administered ripasudil to mice by intraperitoneal (i.p.) injection at 30 mg/kg. Their brains and blood were harvested at 0.5, 1, 2, 4, 8, and 24 hours and processed, and the brain lysates and plasma were used by a mass spectrometer to determine the drug level of ripasudil at each time point. We found fast penetration of ripasudil into the brains, with maximum concentrations reached above 10 µM between 0.5 and 1 hours. Levels of ripasudil rapidly decreased after one hour, and small traces of ripasudil were detected at 24 hours (Fig. 1).

**Figure 1:**
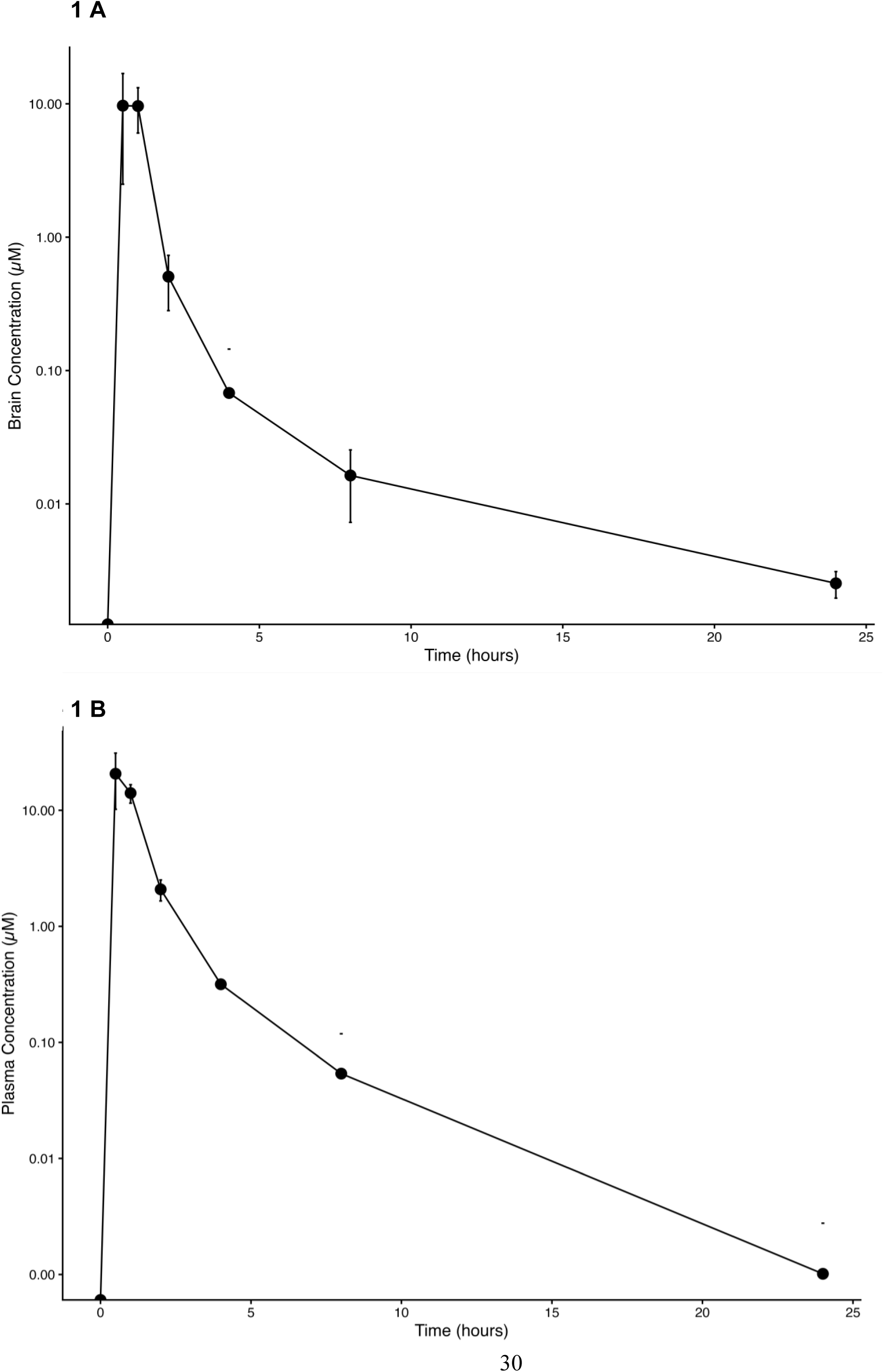
Drug level of ripasudil in mouse brains and plasma. Mice were administered ripasudil by intraperitoneal (i.p.) injection at 30 mg/kg. Mice were sacrificed after 0.5, 1, 2, 4, 8, and 24 hours, and their brains and blood were harvested. The brains and blood were processed, and the lysates and plasma were used by a mass spectrometer to determine the drug level of ripasudil at each time point. A) Brain and B) plasma reached the maximum concentration of ripasudil at 10 µM, which occurred between 0.5 and 1 hours. Levels of ripasudil rapidly decreased after one hour, and small traces of ripasudil were detected at 24 hours. N=2 (1 male, 1 female) for each time point. Data shown is mean±SEM.

### Ripasudil-induced pathway analysis in mouse brain

To understand the impact of ripasudil on molecular pathways that may potentially relate to Tau and mitochondrial function, we performed proteomic profiling of brains from mice treated with ripasudil and compared the profiles to saline-dosed mice. Mice were treated with ripasudil at 30 mg/kg for three hours or saline by i.p. injection before their brains were harvested for proteomic analysis. The selection of time point and dosage was based on the time course and peak brain exposure of ripasudil (Fig. 1). Compared to mice dosed with saline, we identified differentially-expressed proteins in mice dosed with ripasudil and subsequently performed gene ontology enrichment analysis to identify the processes that were significantly affected. In male mice, the seven significantly up-regulated pathways are related to components of chromosomal region, e.g., chromosome, chromosome centromeric core domain, chromosome centromeric region, nucleosome, protein DNA complex, and structural constituent of chromatin (Fig. 2A). In female mice, the significantly down-regulated pathways are related to components of chromosome centromeric core domain and structural constituent of chromatin (Fig. 2B). Pathway network analysis in male mice revealed that the significantly up-regulated pathways are closely related (Fig. 2C). Pathway network analysis mice revealed the significantly down-regulated pathways were the main pathways differentially expressed and closely related (Fig. 2D).

**Figure 2:**
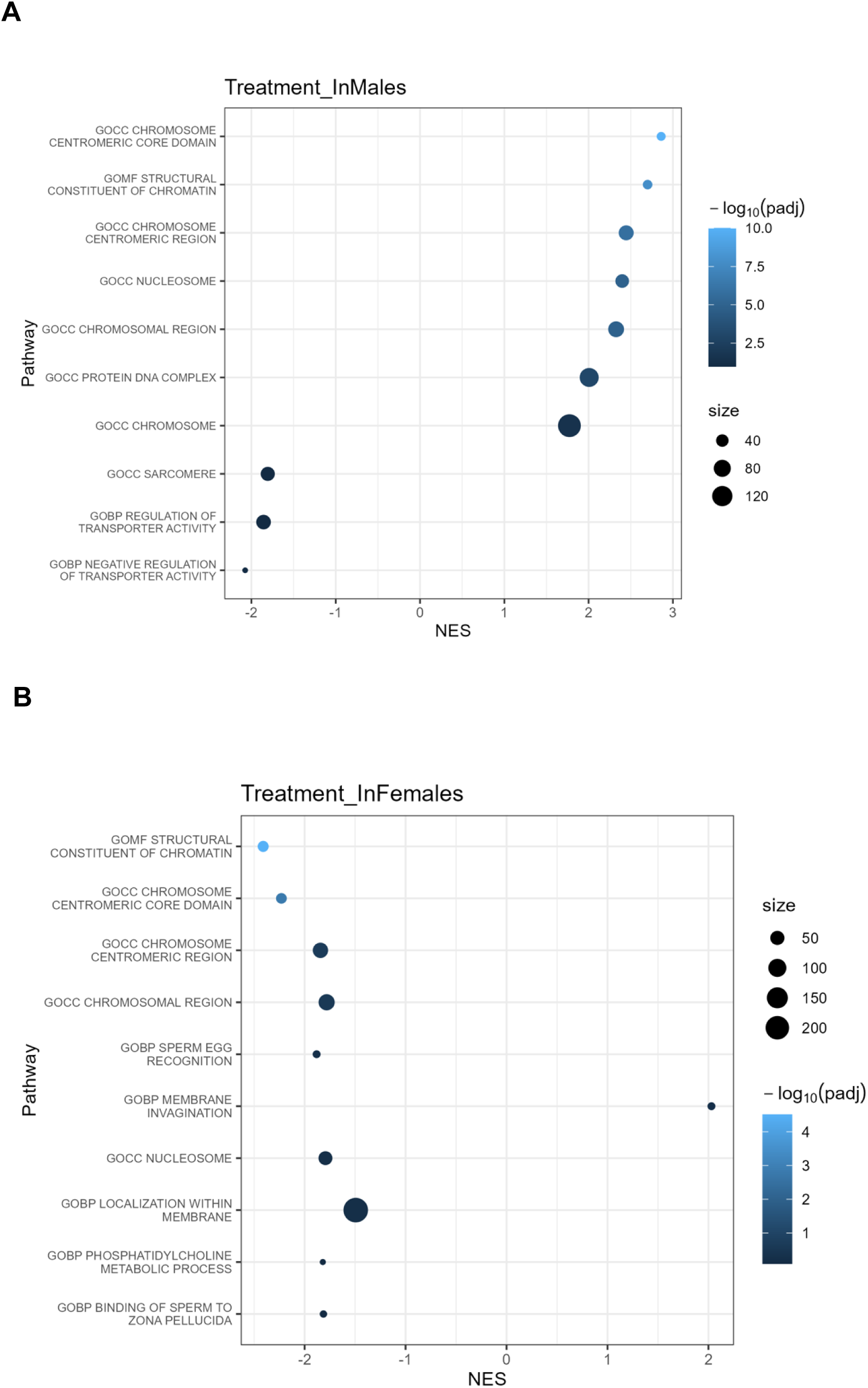
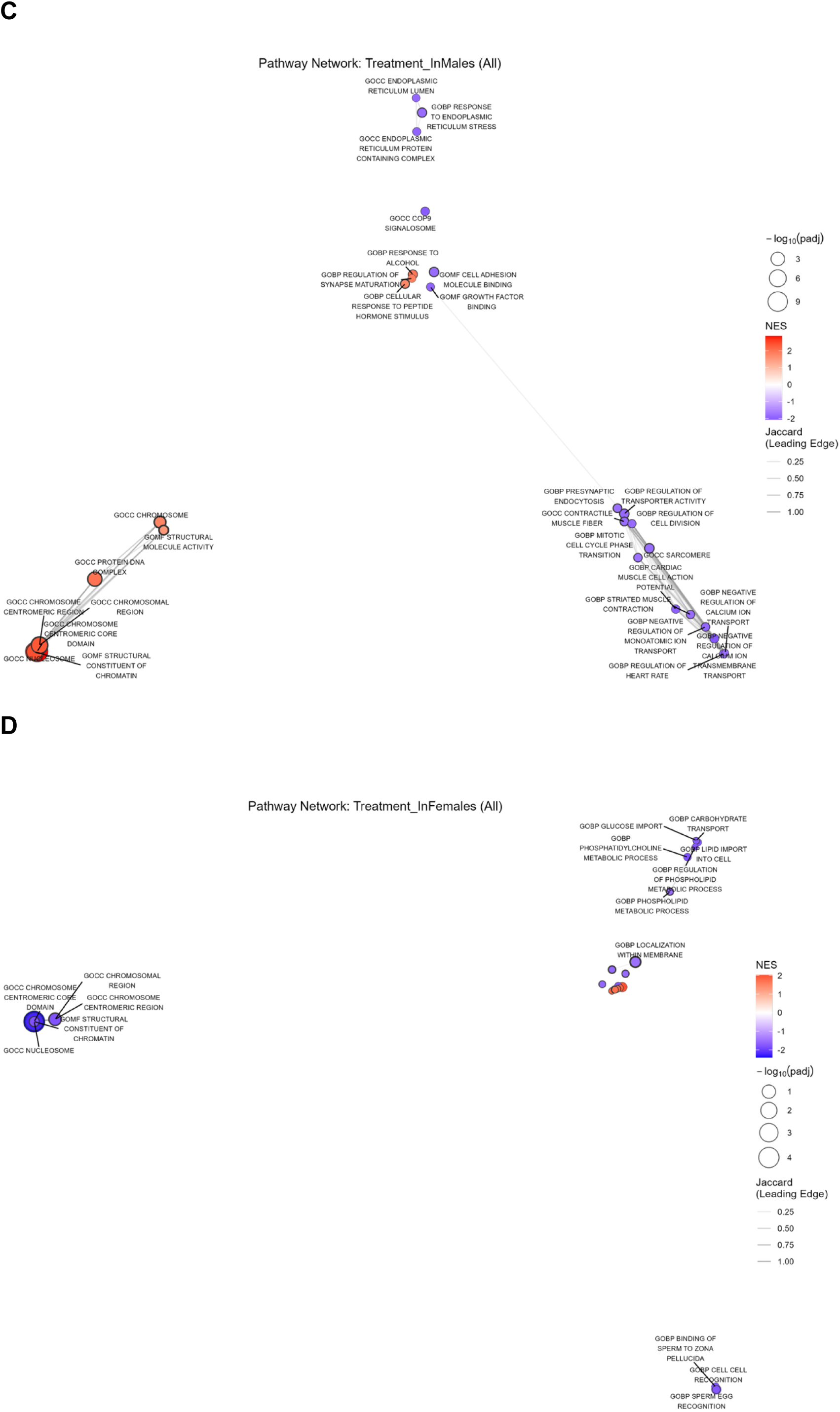
Protein abundance and molecular pathways impacted by ripasudil in mouse brains. A) Up-regulated pathways in male mice and B) Down-regulated pathways in female mice dosed with ripasudil. C and D) Pathway networks were analyzed. Normalized Enrichment Score (NES) was used to denote up-(+) or down-(-) regulation. Size of the points correspond to the -log10(padj) value, with higher value being more significant. Distance between points is roughly the jaccard index of leading edge proteins (the most upregulated proteins) in that pathway. Border thickness around points is reflective of the pathway size. Pathway networks filtered by C) male and D) female were illustrated. N=5 male and N=5 female.

### Ripasudil reduced Tau and pTau 396 in *App^NL-G-F^* knock-in mouse brains

To determine whether ripasudil impacts AD-related proteins *in vivo*, brain lysates were extracted, and Tau and pTau 396 ELISA were performed to measure levels of AD pathological proteins in mouse brains. In *App^NL-G-F^* knock-in mice dosed with ripasudil, there was no significant reduction of Aβ40 in brains, compared to those dosed with saline (Supplemental Fig. 1). Importantly, those mice showed significant reduction of Tau (p<0.0001 for female, p<0.001 for male) (Fig. 3A) and pTau 396 (p<0.05 for female, p<0.01 for male) (Fig. 3B) compared to mice dosed with saline. There is also a significant difference (p<0.001) in Tau between sexes (Fig. 3A).

**Figure 3:**
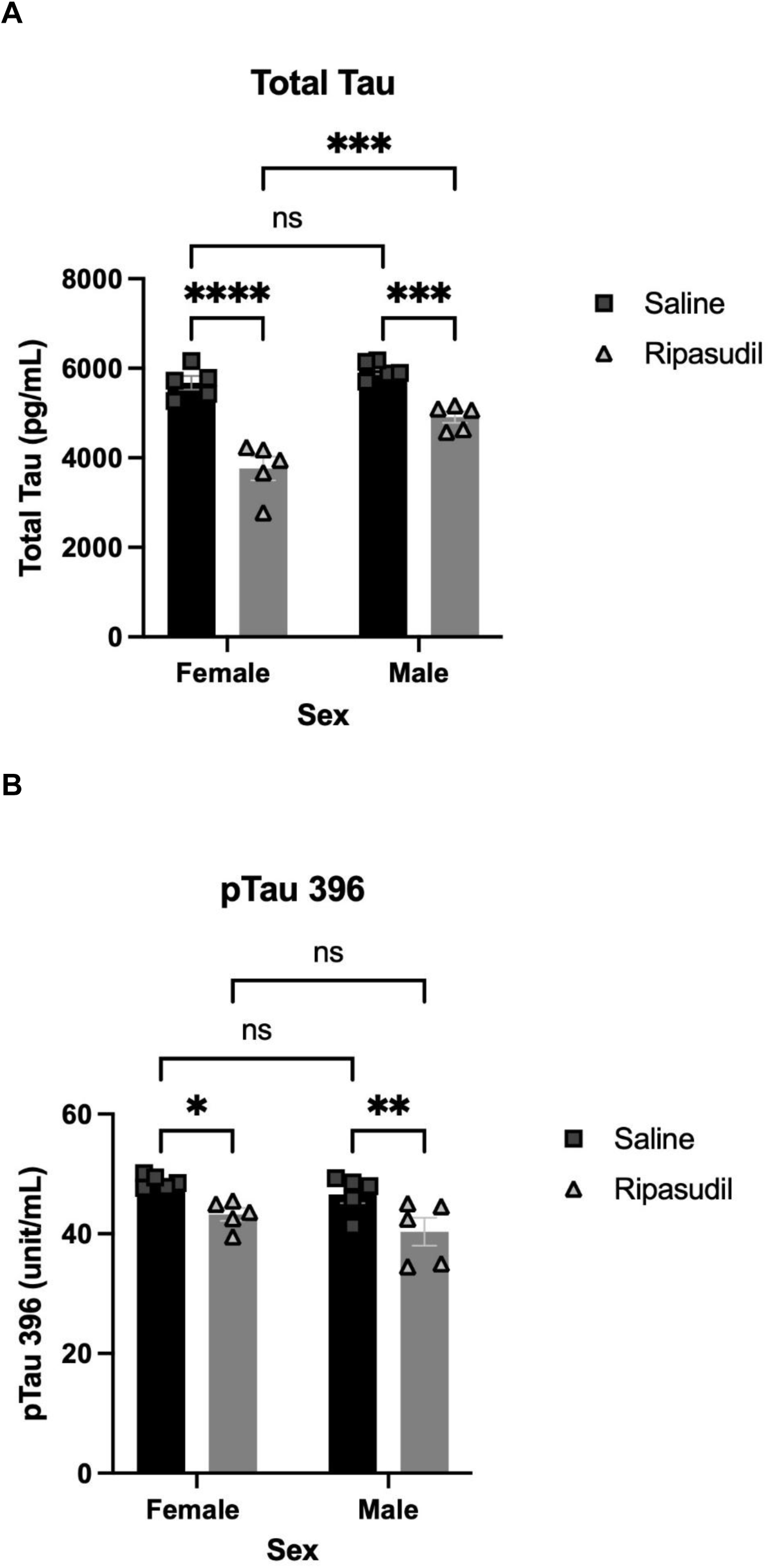
Mouse brains exposed to ripasudil showed significant reduction of Tau and pTau 396. *App^NL-G-F^* knock-in mice were dosed with ripasudil at 30 mg/kg or saline and sacrificed after three hours. Mouse brain lysates were prepared and processed for ELISA (5 male, 5 female *App^NL-G-F^* mice for saline group and 5 male, 5 female *App^NL-G-F^* mice for ripasudil group). A) Ripasudil significantly decreased Tau and B) pTau 396 in *App^NL-G-F^* knock-in mice, compared to saline-dosed mice. Two-way ANOVA and Tukey’s post-hoc tests were used. * denotes p<0.05, ** denotes p<0.01, *** denotes p<0.001, and **** denotes p<0.0001. Data shown is mean±SEM.

### Ripasudil improved mitochondria function

To understand the functional impact of ripasudil on neuronal cells, we cultured neural stem cells derived from AD patients, as previously described by our group ^53, 54^. After we treated iPSC-derived neural stem cells with ripasudil or growth media (control untreated) overnight and three baseline measurements, we measured mitochondrial function, since mitochondrial dysfunction is a common feature involved in AD pathogenesis ^27, 57^. A Mito Stress Test of ripasudil-treated neural stem cells showed 35.2% increases (74.4 vs. 55.01 pmol/min) in basal oxygen consumption (p<0.05), 37.7% (53.83 vs. 39.08 pmol/min) in ATP-linked respiration (p<0.05), 69.3% (172.3 vs. 101.8 pmol/min) in maximal respiration (p<0.01), and 150% (94.48 vs. 37.74 pmol/min) in reserve capacity respiration (p<0.001) compared to untreated cells after overnight treatment (Fig. 4).

**Figure 4:**
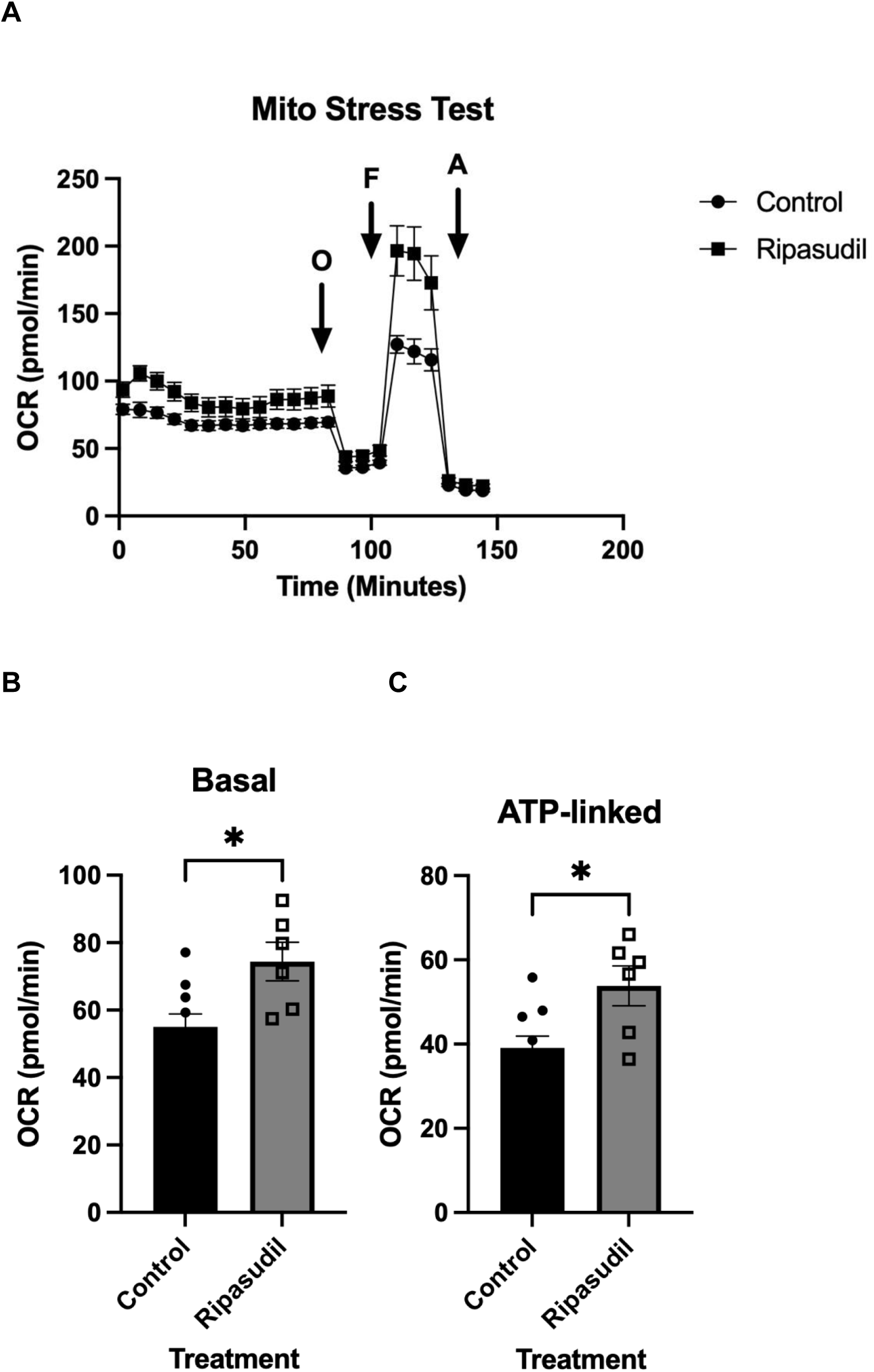
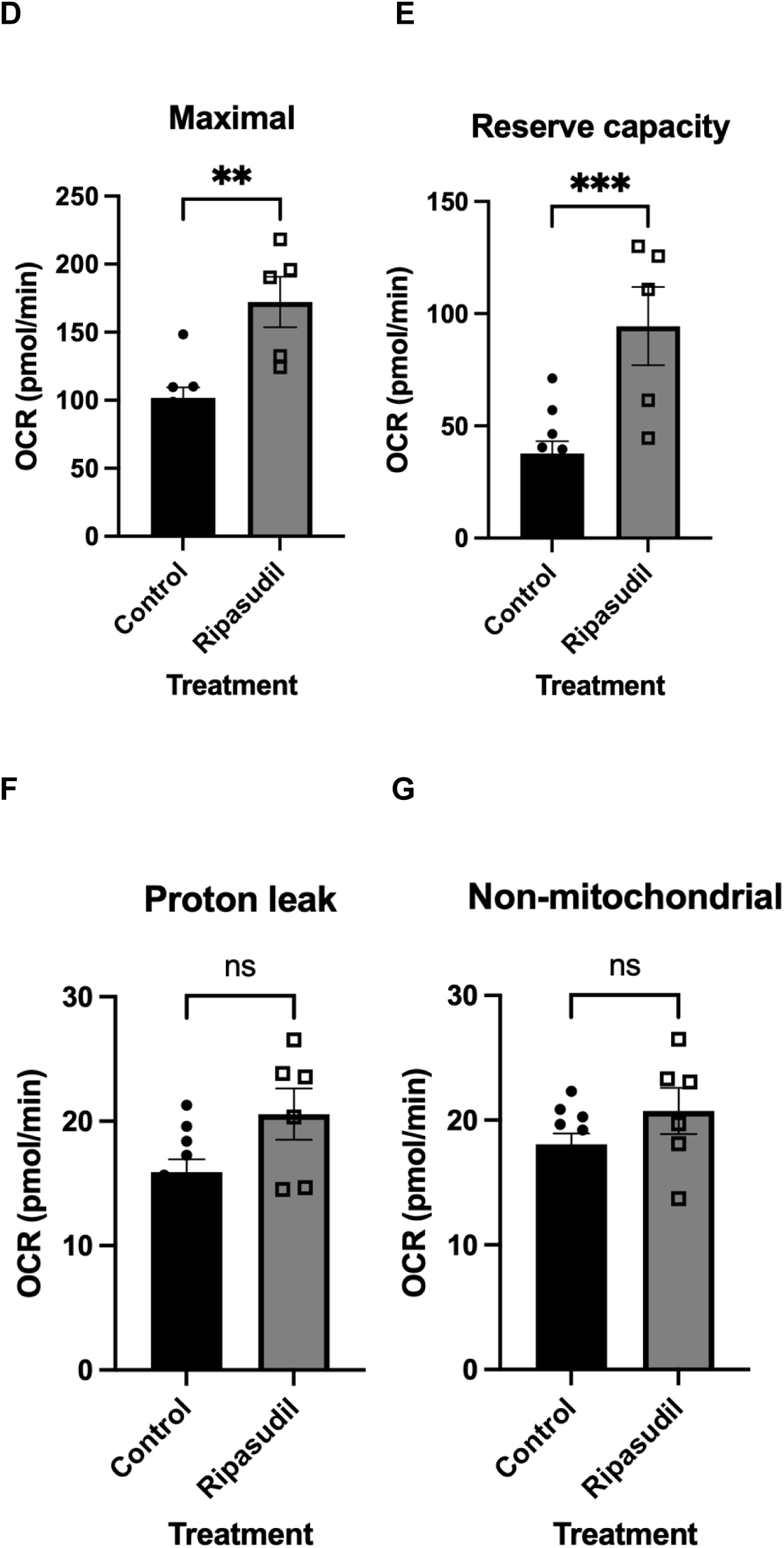
Ripasudil improves mitochondrial function in iPSC-derived neural stem cells. Neural stem cells (NPCs) were plated at 4×10^4^ cells/well the night before the experiment and maintained in the incubator at 37 °C with 5% CO_2_. One hour after cells were seeded (N=10 wells for control, N=6 wells for ripasudil), ripasudil was added at 10 µM in the STEMdiff Neural Progenitor medium for overnight treatment. For the mitochondrial stress test (MST), NPCs were incubated in XF medium with and without ripasudil for 1 hour, and oxygen consumption rate (OCR) was measured continuously for 2 hours at baseline or after injection of mitochondrial inhibitors at indicated time. A) Abbreviation: O = 1.5 µM oligomycin to determine OCR linked to ATP production, F = 2 µM FCCP to determine the maximal OCR, and A = 0.5 µM antimycin A to determine non-mitochondrial OCR. Basal OCR (OCR before O minus OCR after A), ATP-linked (OCR before O minus OCR after O), maximal (OCR after FCCP minus OCR after A), reserve capacity (OCR after FCCP minus basal OCR), proton leak (OCR after O minus OCR after A) and non-mitochondrial OCR (OCR after A) were calculated. Ripasudil significantly increased the oxygen consumption rate for B) basal respiration, C) ATP-linked respiration, D) maximal respiration, and E) reserve capacity respiration. Ripasudil did not increase the oxygen consumption rate for F) proton leak respiration and G) non-mitochondrial respiration. * denotes p<0.05, ** denotes p<0.01, *** denotes p<0.001 compared to control. ns= not significant. Data shown is mean±SEM.

### Acute ripasudil treatment did not increase brain [^18^F]BCPP-EF PET signal in wild-type mice

To explore whether ripasudil impacted mitochondrial function through mitochondrial complex -1 (MC-1) protein *in vivo*, we performed a pilot study of PET imaging using [^18^F]BCPP-EF, a radiotracer designed to image and quantify MC-1 in the brain. Wildtype mice were treated with ripasudil at 30 mg/kg for two hours or saline by *i.p.* injection before PET imaging. The whole brain SUV time-activity curves (TACs) showed lower [^18^F]BCPP-EF uptake in the ripasudil-treated mouse compared to the saline-treated mouse (Supplemental Fig. 2a). Consistent with this observation, cerebellum-normalized SUVR (SUVR_CB_) images showed lower apparent [^18^F]BCPP-EF signal intensity in the ripasudil-treated mouse compared with the saline-treated mouse across the brain regions examined (Supplemental Fig. 2b). These preliminary findings suggest that acute ripasudil administration did not increase [^18^F]BCPP-EF PET signal in the brain under the present experimental conditions.

## DISCUSSION

We applied a conventional small molecule drug development approach to evaluate a drug in clinical use for new application—that is, as a potential new AD therapeutic agent. Instead of creating a large number of derivatives to establish structure and activity relationship of new compounds, we performed pilot studies to explore the feasibility of establishing pathological Aβ and Tau proteins and a mitochondrial functional test as outcomes to evaluate our candidate drug. For this study, we tested ripasudil, a glaucoma drug approved for clinical use in Japan in 2014. We established the time course of ripasudil exposure in mouse brains after a single i.p. injection of 30 mg/kg. The brain penetration of ripasudil was rapid, with maximum concentration observed after 30 minutes, which was similar to results observed in rabbits ^58^. Small traces of ripasudil were observed 24 hours after injection. The rapid brain penetration of ripasudil between 0.5 and 1 hour resulted in a higher concentration (>10 µM) than its IC_50_, which was 0.051 µM and 0.019 µM for ROCK1 and ROCK2, respectively ^35^.

Our proteomic profiling of mouse brains revealed certain pathways activated by ripasudil, without any seeming safety concerns. In male mice, the significantly enriched pathways are related to the components of chromosome, centromere, nucleosome, chromatin, and the protein DNA complex. The DNA-protein complex is an essential assembly that governs how genetic information is packed, repaired, and replicated, and the up-regulation of these ways may suggest ripasudil’s role in transcription and DNA regulation. Our proteomic profiling of mice dosed with ripasudil did not reveal specific pathways related to increases of Tau expression or decreases of Tau degradation.

The most relevant outcome to AD therapeutics is the change of AD pathological proteins such as Tau and pTau, and the acute effect aligning with the improved mitochondrial function, as altered cellular energy states are known to impact pathological processes in AD ^67, 68^. These results are notable given the acute exposure of ripasudil and the use of mice without overexpressing Tau, as significant reductions of Tau and pTau 396 were all detected in *App^NL-G-F^* knock-in mouse brains. Our data suggest that ROCK inhibition and ripasudil may influence early molecular changes that precede plaque and tau tangle formation, further solidifying its candidacy as an AD therapeutic agent.

The most important and consistent feature of ripasudil is that it improved mitochondrial function in neural stem cells, which corroborated our findings in the proteomic study. Because the neural stem cells were derived from AD patients, they are clinically relevant. Ripasudil significantly increased oxygen consumption rates in basal respiration, ATP-linked respiration, maximal respiration, and reserve capacity respiration, indicating an overall enhancement of mitochondrial performance. Increased ATP-linked respiration suggests increased ATP demand, possibly from increased cellular activities, such as oxidative phosphorylation. Given that impaired ATP generation is a well-documented feature of AD ^69, 70^, enhancement of this parameter is particularly relevant. Furthermore, reserve capacity measures cells’ ability to survive stressors such as Ca^2+^ imbalance and reactive oxygen species (ROS). Increased reserve capacity and maximal respiration suggest cells’ improved ability to respond to energetic changes. These findings align closely with our proteomic data, which demonstrated the enrichment of pathways involved in mitochondrial function and energy generation. Together, these results suggest that ROCK inhibition through ripasudil engages in the regulation of mitochondrial function and efficiency, further supporting ripasudil’s neuroprotective effects in the context of AD.

In this pilot PET imaging experiment, acute ripasudil treatment did not increase brain [^18^F]BCPP-EF PET signal in wild-type mice. Whole-brain SUV showed lower early tracer uptake in the ripasudil-treated mouse compared with the saline-treated mouse, although uptake became more comparable during the later phase of image acquisition. Consistently, cerebellum-normalized SUVR images showed lower apparent [^18^F]BCPP-EF signal intensity after ripasudil treatment. These findings should be interpreted cautiously because of the pilot nature of the experiment and the acute 2 h treatment paradigm. The PET signal may also be influenced by tracer delivery, tissue kinetics, radiotracer metabolism, and regional brain physiology. Therefore, the present data suggests that acute ripasudil administration did not produce a detectable increase in MC-1-related PET signal under the conditions tested. Longer treatment paradigms, larger sample sizes, and complementary *ex vivo* validation will be required to determine whether ripasudil modulates mitochondrial function or MC-1 expression in the brain.

Overall, our results provide evidence that ripasudil improved mitochondria function, and reduced Tau and pTau levels in mouse brains. Because ripasudil has already been in clinical use in Japan, repurposing ripasudil may reflect a faster path than the conventional drug development process. Therefore, it would potentially be on a fast-track for FDA evaluation and approval as an AD therapeutic agent.

## Acknowledgment

This work was supported in part by UAB Nathan Shock Center P30 AG050886 (VDU, JZ), and NIH R01 AG063913 (WX) and the Department of Veterans Affairs I01 BX004730 (WX), and the Canadian Institutes of Health Research PJT191769 (CZ). JP and CZ thank Centre for Addiction and Mental Health BHIC staff for their professional technical assistance in the cyclotron operation and PET imaging studies. The views expressed in this article are those of the authors and do not represent the views of the US Department of Veterans Affairs or the US Government.

Declaration of Generative AI and AI-assisted technologies in the writing process.

AI was not used.

## Data Availability

Data is available upon submitting request to the corresponding author.

## Ethical approval

All studies were approved by the institutional IACUC at the Bedford VA Healthcare System.

## Supplemental material

**Figure 1:**
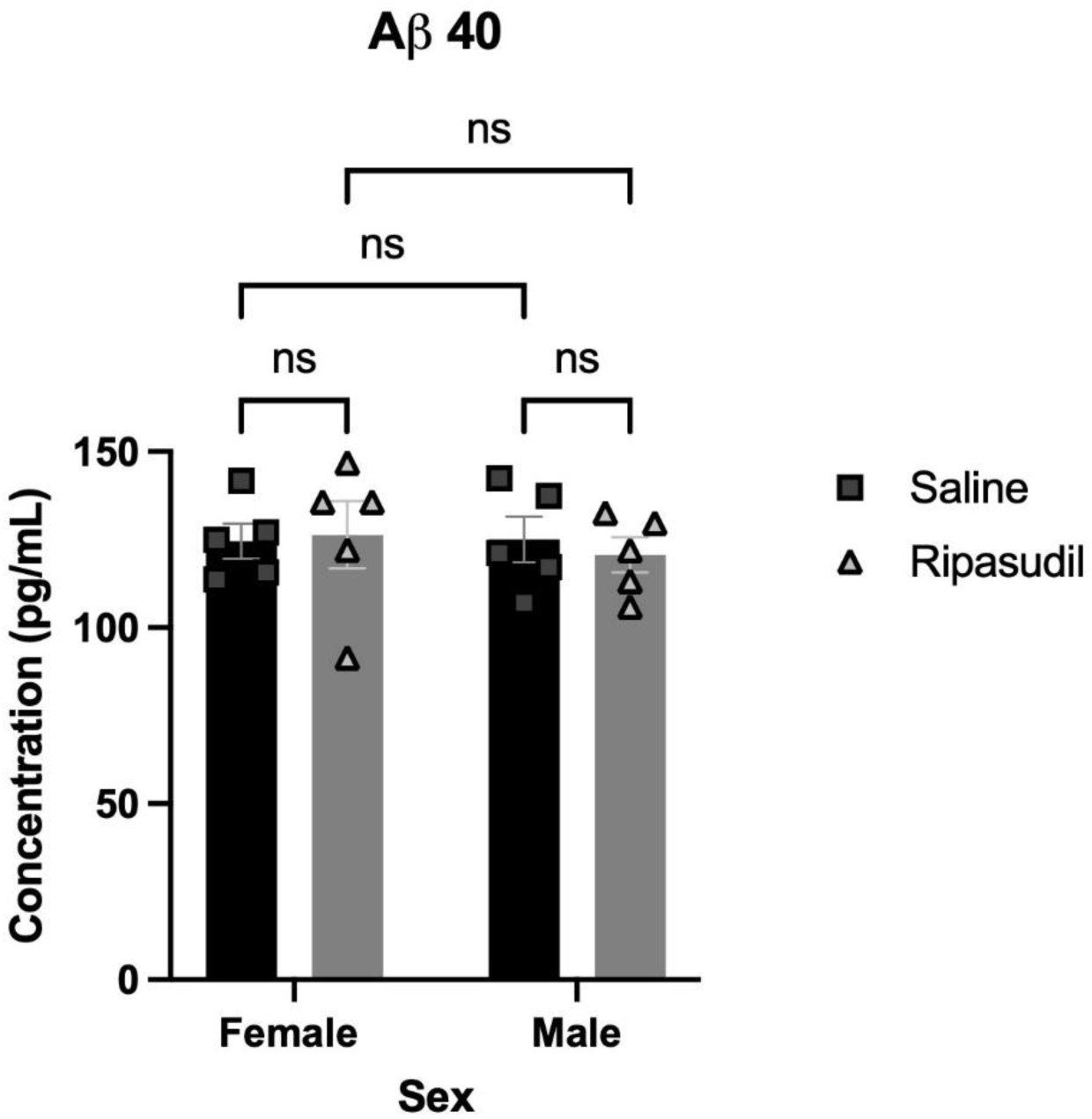
*App^NL-G-F^* knock-in mouse brains exposed to ripasudil did not show significant reduction of Aβ. Mice were dosed with ripasudil at 30 mg/kg or saline and sacrificed after three hours. Mouse brain lysates were prepared and processed for ELISA (5 male, 5 female mice for either ripasudil or saline group). Ripasudil did not significantly decrease Aβ40 compared to saline-dosed mice. Data shown is mean±SEM.

**Figure 2:**
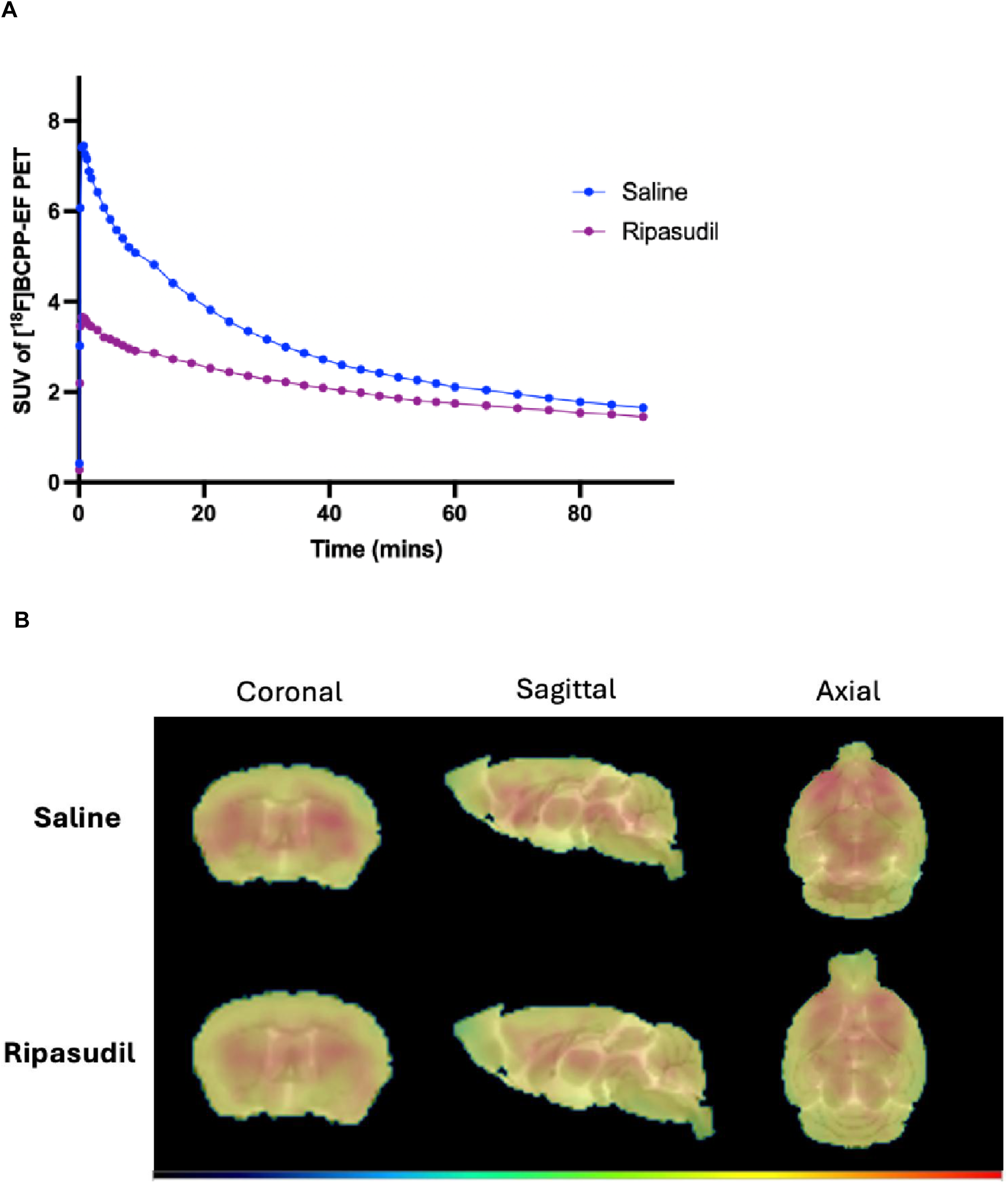
Acute ripasudil treatment did not increase brain [^18^F]BCPP-EF PET signal in wild-type mice. Six-month-old female wild-type C57BL/6 mice were administered either saline or ripasudil (30 mg/kg) i.p. 2 hours prior to [^18^F]BCPP-EF PET/CT imaging. (A) Whole-brain SUV time–activity curves of [^18^F]BCPP-EF uptake in the ripasudil-treated mouse and the saline-treated mouse. (B) Representative cerebellum-normalized SUVR PET images of [^18^F]BCPP-EF signal intensity in the ripasudil-treated mouse compared with the saline-treated mouse. These pilot data suggest that acute ripasudil treatment did not increase mitochondrial complex I-related PET signal under the present experimental conditions.

## Notes

### Competing Interest Statement

The authors have declared no competing interest.

